# Intracortical vessel density mapping at mesoscale reveals regionally specific angioarchitecture in the human brain

**DOI:** 10.64898/2026.05.01.722251

**Authors:** O.F. Gulban, K. Wagstyl, L. Huber, A. Pizzuti, Saskia Bollmann, A. Roebroeck, R. Goebel, K. Kay

## Abstract

The metabolic demands of the human brain are met by a complex vascular architecture, yet our characterization of this network remains incomplete. While we have mapped the macroscopic vessels on the brain’s surface and the microscopic capillaries within small tissue samples, the *mesoscopic scale* consisting of the penetrating vessels that plunge through cortex remains an anatomical terra incognita. Mapping the interface between the macroscopic and microscopic scales is essential to understanding the critical vascular supply that sustains brain health. Here, we leveraged the BigBrain dataset and developed custom detection and tracing algorithms to reveal a whole-cortex record of the mesoscopic vascular network. We find that vascular density is not uniform across the cortex, but is a heterogeneous landscape that shows clear relationships to traditional areal boundaries. Our results constitute a reference for human mesoscopic angioarchitecture and demonstrates the power of repurposing high-resolution histological atlases. Ultimately, this work lays the groundwork for validating recently developed in vivo MRI techniques for imaging the human cerebrovascular system at mesoscale.

## 1 Introduction

The human brain relies on a hierarchical vascular architecture to sustain its immense metabolic demands (Drew, 2022; Polimeni, 2025). Traditionally, neuroanatomical models have treated the penetrating arterioles and venules (in short: *meso-vessels*) that bridge macroscopic surface delivery and microscopic capillary perfusion as a homogeneous, unpatterned backdrop (Duvernoy et al., 1981). Under this assumption, the meso-vessels are viewed as a uniform structure rather than a specialized system that varies across the cortex. This has led to a reliance on generalized average metrics, such as the reported 2.2-to-1 ratio of intracortical arteries to veins in humans (Schmid et al., 2019). However, such global ratios are often derived from localized samples and may not accurately capture regional heterogeneity across regions. Furthermore, even if a ratio remains constant, the absolute density of vessels across the cortex can vary (see **Figure 1**).

**Figure 1:**
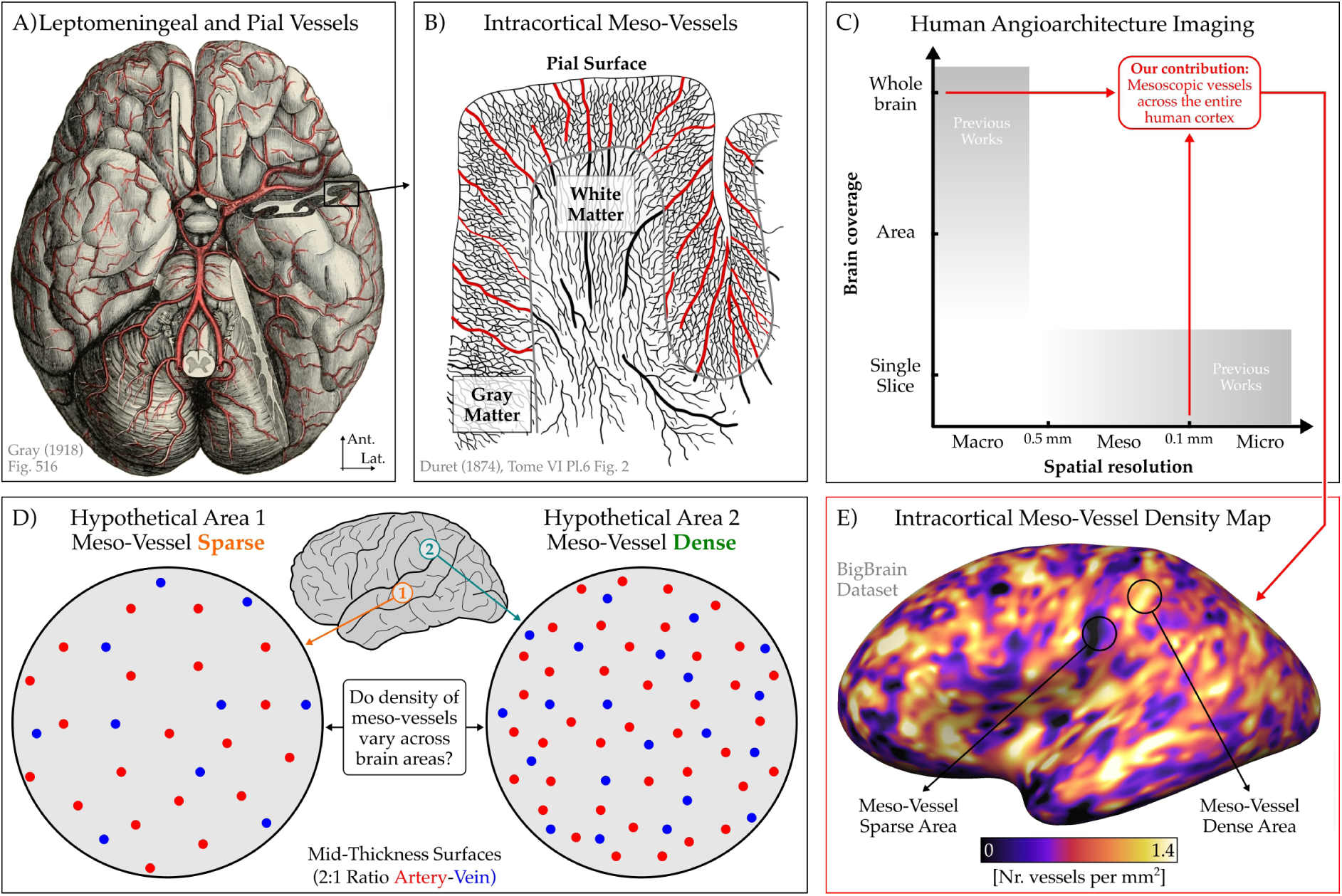
Vascular nomenclature and the gap in understanding the vasculature at mesoscopic scale. **Panel A** shows macroscopic angioarchitecture of the human brain (only arteries are depicted for clarity). These surface vessels are further classified into leptomeningeal and smaller pial vessels according to Duvernoy and Vannson (1999). **Panel B** shows intracortical mesoscopic vessels. Highlighted in red are the radial meso-vessels (which are also called descending arteries and ascending veins (Schmid et al., 2019) while smaller lateral branches are shown alongside the capillary bed in black. **Panel C** shows coverage and resolution of current efforts to measure human cerebral vasculature. Current imaging efforts are bifurcated between macroscopic surface views and microscopic capillary reconstructions. Macroscopic imaging lacks the resolution to resolve intracortical structures, while microscopic histology lacks the whole-brain coverage required to identify regional variations. This leaves a critical knowledge gap at the mesoscopic scale (0.1 to 0.5 mm). **Panel D** shows our research objective: investigating whether the spatial density of meso-vessels varies systematically across cortical areas. **Panel E** shows our primary contribution: the first comprehensive, whole human brain mesoscopic vessel density map, bridging the gap between macroscopic and microscopic vascular anatomy. The drawing in Panel A is adapted from Gray and Lewis (1918) Figure 516 (public domain) and the drawing Panel B is adapted from Duret (1874) Tome VI Pl. 6 Figure 2 (public domain).

Whether the density of intracortical meso-vessels is homogenous or varies heterogenously across regions is a fundamental open question of brain organization. Foundational ex vivo work in primates originally demonstrated that microvascular density is localized, showing discrimination between the primary visual cortex and surrounding areas (Pfeifer, 1940), alongside elevated capillary density within metabolic cytochrome oxidase blobs (Zheng et al., 1991). Recent evidence in macaques suggests that there is substantial areal heterogeneity (Autio et al., 2024, 2025; Wang et al., 2025). However, a similar map has not been measured in humans at mesoscale (0.1 to 0.5 mm). Therefore, we do not yet know: (I) whether the meso-vessels are distributed uniformly, (II) whether they co-vary with cortical geometry (thickness, curvature), or (III) whether they vary with the specialized metabolic requirements of local neural populations in humans (see Weber et al. (2008) for macaques). Determining the degree with which meso-vessel density varies is essential to understanding whether the intracortical vascular network possesses a distinct architectural logic.

We demonstrate in this paper that the mesoscopic vascular network is not a uniform grid, but a highly organized system characterized by substantial regional heterogeneity. Using the BigBrain dataset (Amunts et al., 2013), which is a high-resolution 3D digital reconstruction of a human brain made out of 7400 histological sections at 20 micron thickness with cell body staining, we discovered an untapped record of the intracortical meso-vessels. By applying tailored vessel enhancement and detection algorithms, we transformed this cell-body atlas into a comprehensive map of cortical meso-vessel density for an entire human brain (see **Figure 2**).

**Figure 2:**
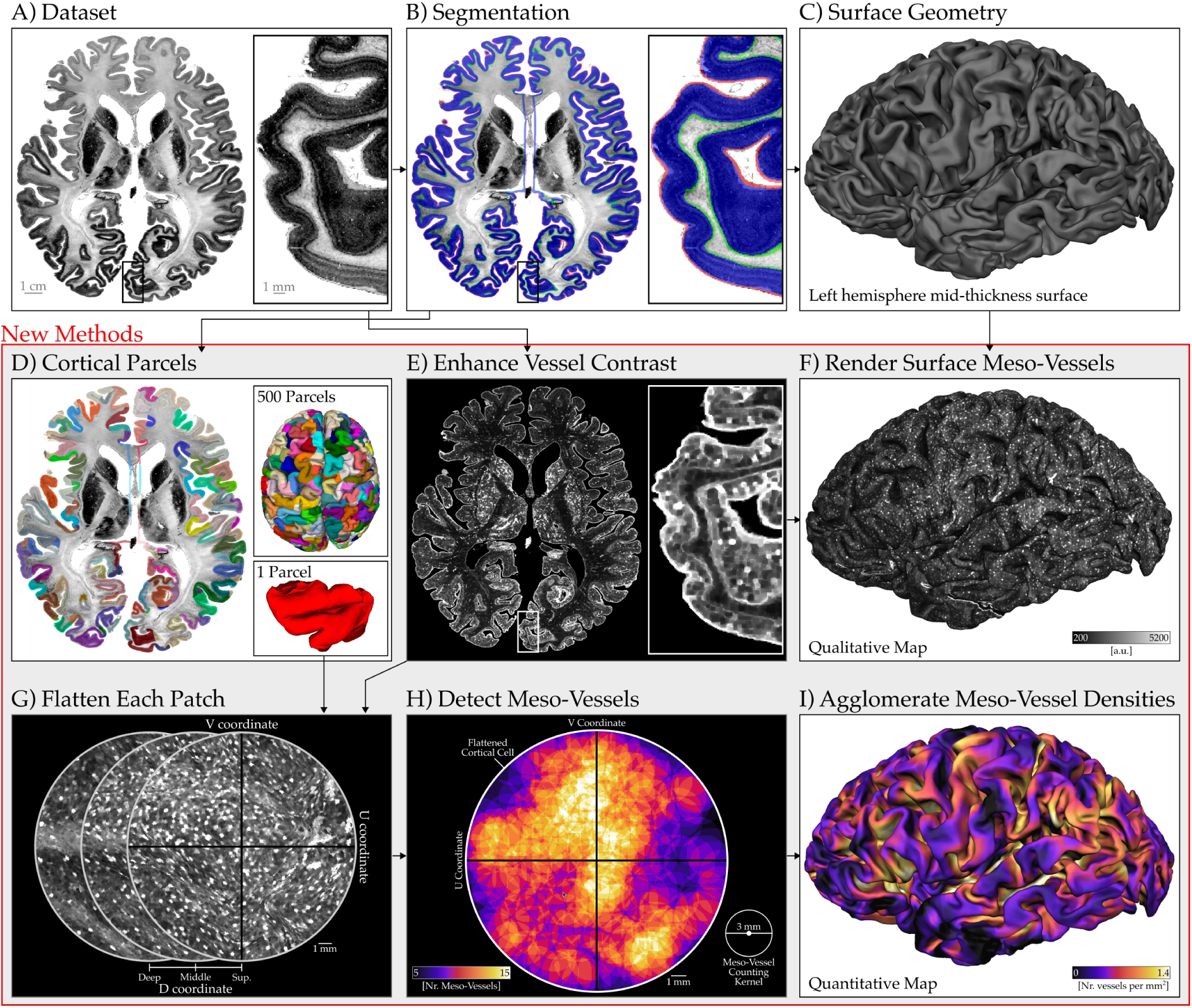
Methods overview for meso-vessel density mapping. The term vessel is used here as a collective category for both arteries and veins, as the image contrast does not support the unambiguous differentiation of arteries and veins. **Panel A** shows a high-resolution histological slice of the initial 100 micron isotropic BigBrain dataset. **Panel B** shows our cortical segmentation (red: outer cortical surface, blue: gray matter, green: inner cortical surface). **Panel C** shows our cortical surface reconstruction. The mid-thickness triangular mesh of the left hemisphere (12,923,304 vertices) provides the geometric scaffold for visualizing intracortical meso-vessel details. **Panel D** shows our geometric cortical parcellation. To facilitate high-throughput analysis, the cortex was partitioned into discrete patches for parallelized processing of the massive 100 micron isotropic whole human brain dataset. **Panel E** shows our vascular contrast enhancement. A custom filter was applied to the histological volume to isolate and amplify subtle vascular signatures (white dots) from the surrounding parenchyma. **Panel F** shows the vessel-enhanced volume sampled onto the mid-thickness surface triangular mesh. **Panel G** shows depth-wise flat-patch projection. Penetrating vessels are visualized across multiple cortical depths. This flattened representation reveals the vertical continuity of the mesoscopic network. **Panel H** shows meso-vessel detection calculated within overlapping circular windows of diameter 3 mm. **Panel I** shows the agglomerated meso-vessel density maps which are computed by dividing the number of meso-vessels in each counting kernel with the kernel area multiplied by the areal shrinkage correction factor. These meso-vessel density map is the main result and novelty of our paper, where regional variations can be traced across the cortical landscape.

Our results reveal that vascular density varies systematically across the cortex, providing a new structural feature for defining cortical areas. This discovery implies that the mesoscopic vasculature is not a passive hemodynamic conduit, but rather an active, specialized component of cortical organization. Our findings suggest that the human vascular system might possess an independent architectural logic at the mesoscale. We make our maps freely available at https://doi.org/10.5281/zenodo.19954660, allowing future studies to better understand how the vasculature may relate to other structural and functional properties of the brain.

## 2 Results

### 2.1 Heterogeneity of the intracortical meso-vessel landscape

To bridge the gap between microscopic histology and mesoscopic anatomy, we developed a high-throughput pipeline that partitions the BigBrain hemispheres into computationally manageable “chunks” before projecting them into a flattened coordinate system. Within these flattened patches, we applied an automated tracing algorithm that identifies radial meso-vessels based on their vertical persistence through the cortical layers, ultimately restitching these localized detections into a unified, 3D whole-brain manifold (see **Figure 2 Panel D-I**). Note that we use the term vessels as opposed to arteries or veins because we cannot definitely specify whether they are arteries or veins.

This reconstruction yields a comprehensive map of radial meso-vessels across the entire human cortex. Our analysis shows that meso-vessel density is spatially variable rather than uniform (see **Table 1** and **Figure 3**), with values ranging from 0 to around 2.1 radial meso-vessels/mm^2^. Importantly, the lowest density values were consistently observed in regions containing tears (see **Figure 3** and **Supplementary Figure 1**). The absence of detections in these artifact-prone zones indicates the specificity of the algorithm and confirms that white gaps in the BigBrain data caused by tissue damage were successfully distinguished from the vascular features.

**Figure 3:**
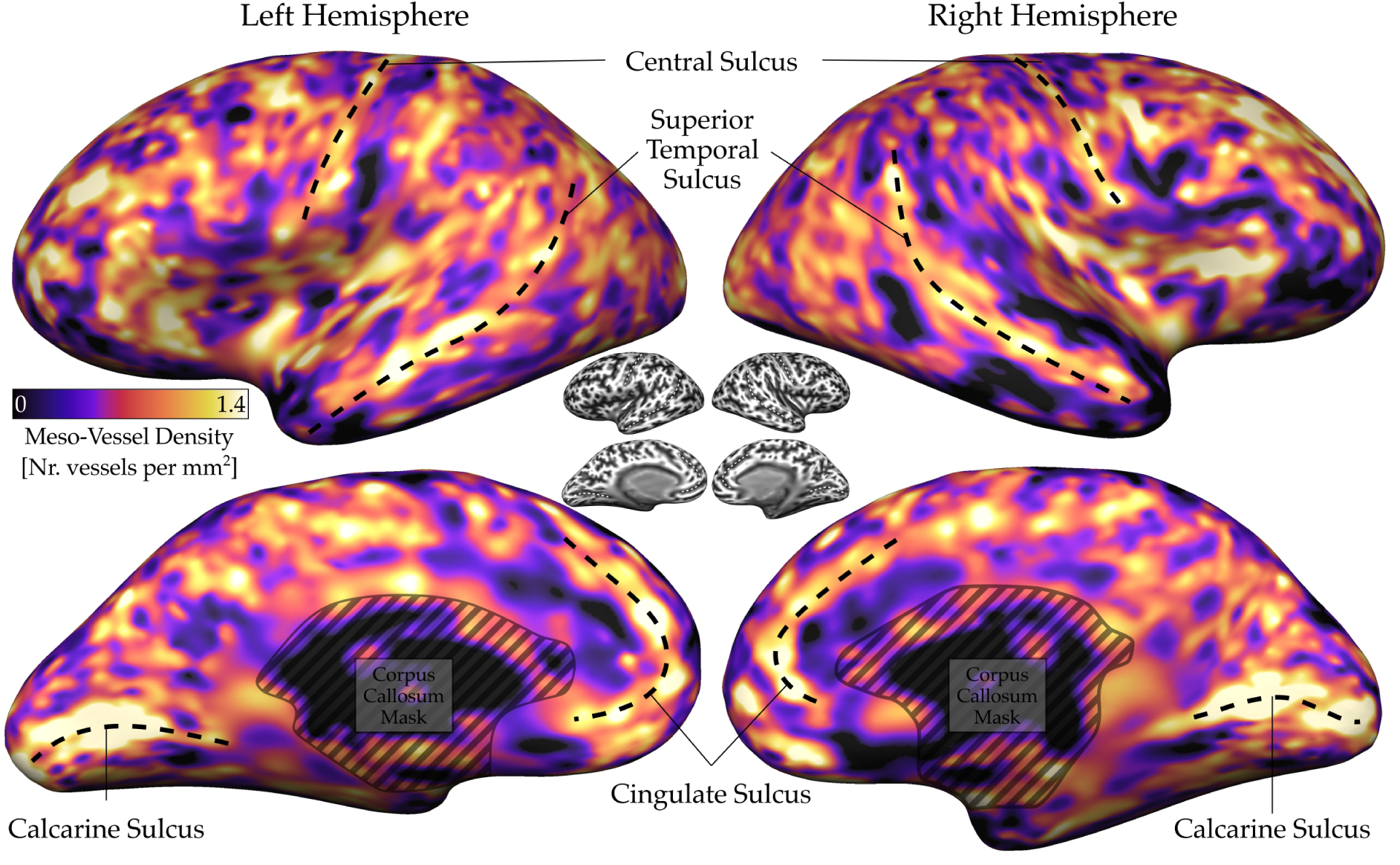
Regional heterogeneity of cortical meso-vessel density. We define meso-vessel density as the number of penetrating vessels per square centimeter of the cortical surface. Our maps reveal that the vascular supply is not uniform. The highest densities are concentrated in the primary sensory and somatosensory regions, suggesting a link between vascular infrastructure and highly metabolically demanding neural computation at the primary areas. The insets show curvature maps where the same sulci are also highlighted.

**Table 1:** Mean meso-vessel density across anatomical lobe. This table summarizes the average meso-vessel density for each major cortical lobe. While these aggregate measures suggest a high degree of global uniformity across large-scale regions, they mask the local heterogeneity visible in **Figures 3–4**. These averages should therefore be interpreted as baseline values that do not reflect the granular mesoscopic angioarchitectural differences visible across the cortical areas.

|  | Left Hemisphere |  | Right Hemisphere |  |
| --- | --- | --- | --- | --- |
|  | Mean | Std | Mean | Std |
| Frontal | 0.719 | 0.195 | 0.717 | 0.215 |
| Occipital | 0.761 | 0.198 | 0.733 | 0.187 |
| Parietal | 0.723 | 0.163 | 0.706 | 0.179 |
| Temporal | 0.664 | 0.198 | 0.604 | 0.203 |
| All Lobes | 0.718 | 0.193 | 0.696 | 0.205 |

The distribution of penetrating vessels across the human cortex reveals a landscape of high-density and sparse areas (**Figure 3**). When quantified as the number of vessels per mm^2^, the vascular supply does not appear to be uniform, but rather variable in nature. The highest concentrations of vessels are found in the primary visual and somatosensory regions (defined based on the atlas provided by Glasser et al. (2016) (see **Figure 4**). Specifically, the primary visual area exhibits a dense, intricate lattice of penetrating vessels, far exceeding the density found in neighboring association zones.

**Figure 4:**
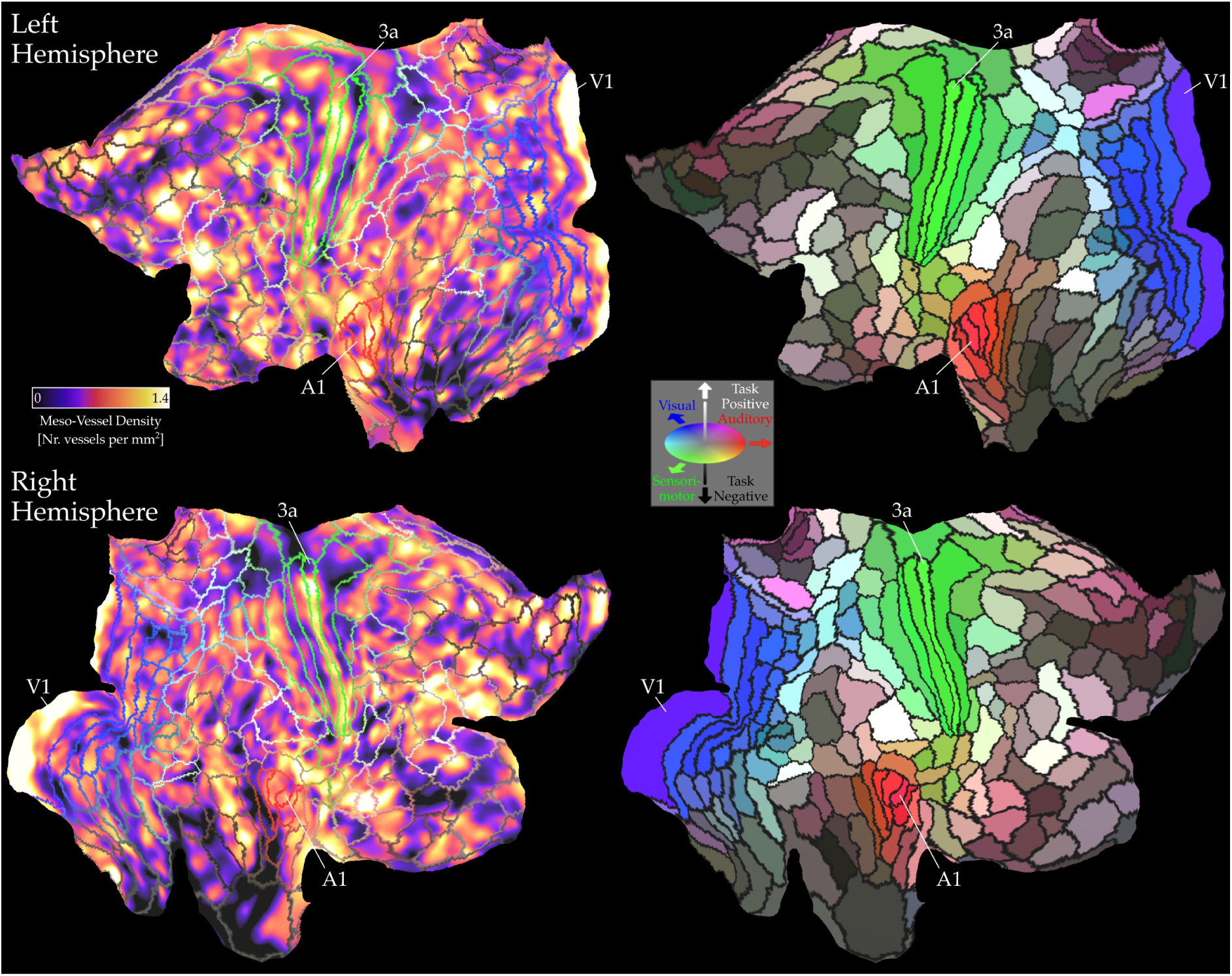
Meso-vessel density map projected onto flattened cortical surfaces of the HCP’s multimodal cortical parcellation (HCP MMP1.0). Primary sensory areas are labeled for reference. It can be seen that the areas V1 and 3a show high meso-vessel density. See **Supplementary Figures 2-4** for curvature, thickness, and mean gray value (MGV) of the Merker stained slices.

In the visual and somatosensory cortex, the vascular network reaches a peak density that suggests a tight coupling between vascular infrastructure and the heavy metabolic demands of primary sensory processing. This regional intensification suggests that the mesoscopic vasculature may be co-developed with the specialized neural architecture of these areas. Specifically, the high vessel density in primary visual cortex may support the fine-grained population receptive field properties required for high-acuity retinotopy, while the enrichment in the postcentral gyrus may align with the high-resolution somatotopic organization of the sensory homunculus.

### 2.2 Divergence of meso-vessel density and cell density

To determine how the meso-vessel density aligns with established cortical landmarks, we projected the meso-vessel density maps onto a cortical surface positioned at the middle of the gray matter and performed correlation analysis against cortical thickness, cortical curvature, and the mean gray value of the Merker stained slices (see **Figure 5**). This comparison reveals that while the vascular network follows its own distinct spatial logic, it remains tethered to the cortical geometry while being distinct from the regional density of cell bodies.

**Figure 5:**
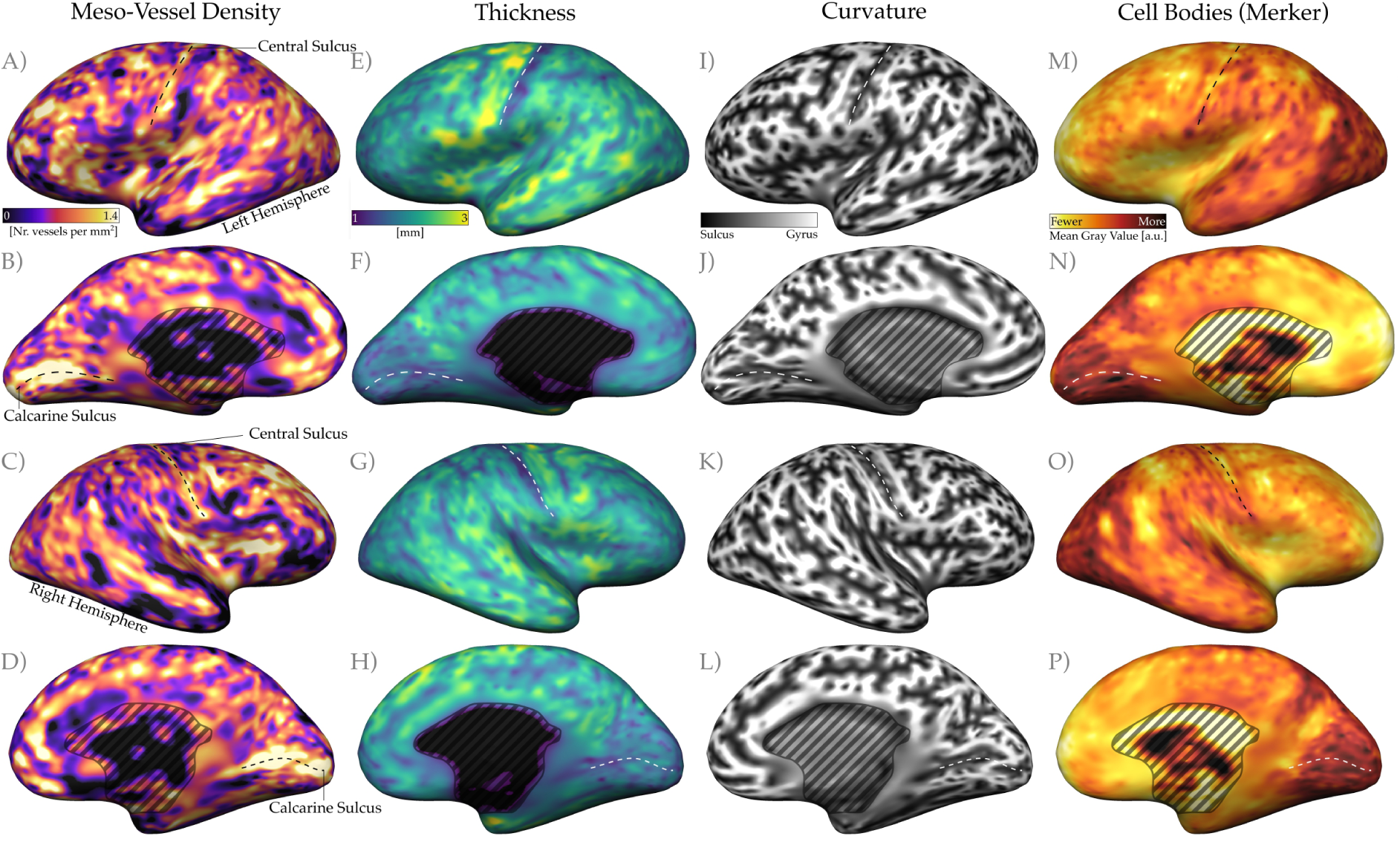
Comparison of meso-vessel density to other anatomical measures. To determine if vascular architecture respects traditional neuroanatomical boundaries, we compared our meso-vessel density maps with established cortical markers: cortical thickness, curvature, and cell-body density. All results are plotted on an inflated mid-gray cortical surface. The cell bodies map was derived from the mean gray values (MGV) of the Merker-stained histological sections. Therefore, darker colors represent higher cell-body density, reflecting the reduced light transmission characteristic of densely packed somata in the BigBrain sections.

One of the most salient features in the meso-vessel density maps is a high plateau of vessel density within the calcarine sulcus (**Figure 3-4)**, which presumably is related to the intense metabolic demands of the primary visual cortex (V1). As one moves toward the gyral banks bordering the sulcus, this density falls away sharply (this is consistent with Hildebrand et al. (2024) Preprint v1 Figure 5). This abrupt transition is consistent across both hemispheres and mirrors the known V1-V2 boundary, marking a clear property of the vascular network. Another feature of the meso-vessel density maps is a high degree of hemispheric symmetry, providing a strong indication that our quantification of meso-vessel density is highly robust and not substantially corrupted by measurement noise.

### 2.3 Meso-vessel density in relation to other cortical properties

To quantify the relationship between cortical geometry and its angioarchitecture, we performed a vertex-wise correlation between the meso-vessel density maps and the geometric properties of the cortical mesh. The distribution of mesoscopic vessels exhibits a consistent but modest relationship with traditional cortical metrics. We found that cortical curvature (negative values indicate sulci and positives indicate gyri) serves as the strongest predictor of vessel distribution, yielding a moderate correlation of r=-0.37 in both hemispheres. The sulci tend to be more meso-vessel dense. This suggests that the physical folds of the brain are associated with the density of the meso-vessels that penetrate through them.

To evaluate the reliability of these correlations, we repeated our analysis across each cortical lobe in each hemisphere (**Table 2**). The results reveal a remarkably stable set of correlations that persist regardless of regional anatomy. Across the frontal, parietal, temporal, and occipital lobes, curvature remained the most robust predictor of vessel distribution, consistently outperforming cortical thickness and cell-body density. The curvature is followed by the thickness, which showed r=-0.26 in left r=-0.33 in the right hemisphere across all lobes. Meso-vessel density versus the mean gray value (Merker) of the histological images showed very weak correlations (r=-0.08 in both hemispheres across all lobes). Therefore, the general pattern of meso-vessel dense sulci and meso-vessel sparse gyri suggests that the mesoscopic vasculature is not a patchwork of local adaptations, but has consistent architectural features. The density of penetrating vessels appears to be governed to some degree by the broad geometric constraints of the cortical ribbon (such as its folding and its thickness) rather than the average cell body density across the brain.

**Table 2:**
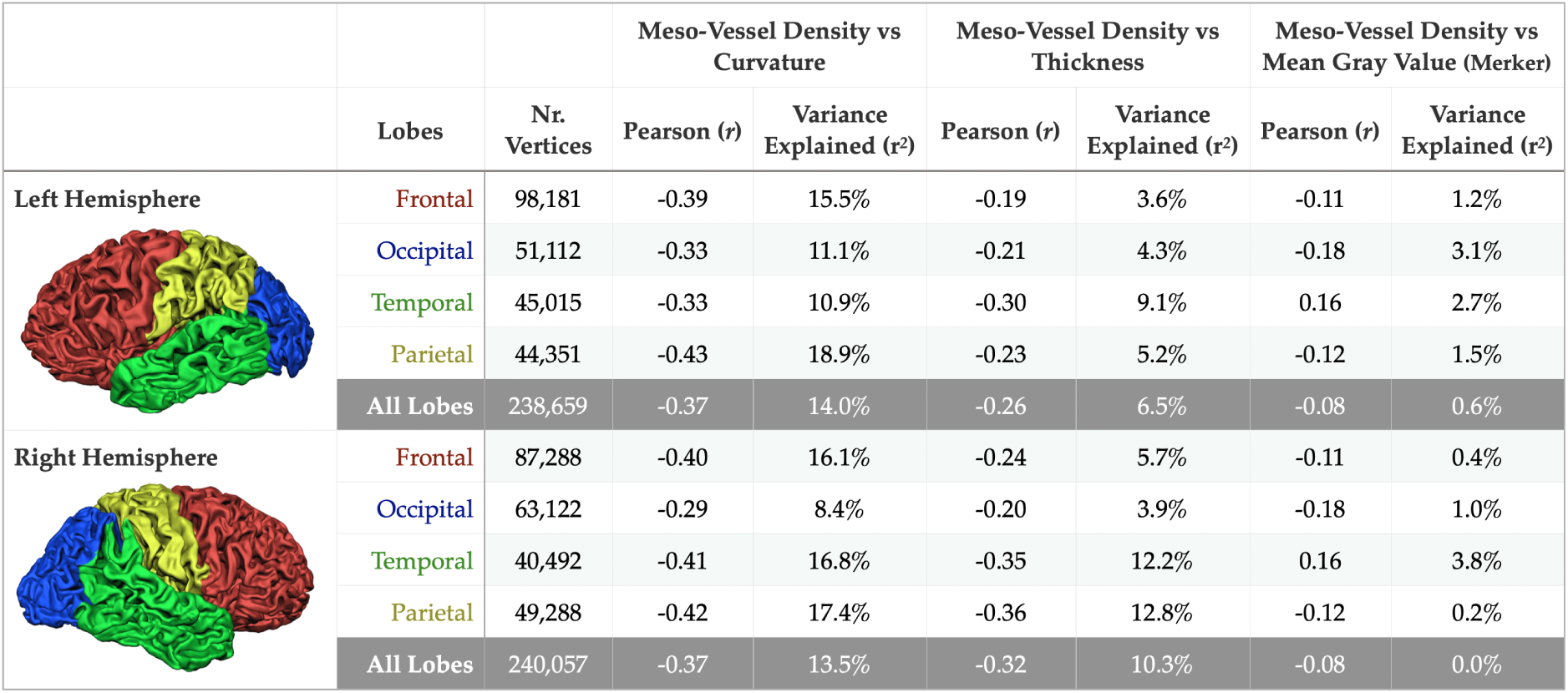
Correlation between meso-vessel density and other cortical measurements. This table summarizes the relationship between meso-vessel density and three key morphometric and histological measures: cortical curvature, thickness, and Merker-stained mean gray values (a proxy for cell-body density). Curvature and thickness were evaluated as potential geometric constraints on the angioarchitecture. For instance, the occupation of sulcal fundi by large pial vessels may influence the distribution and branching patterns of the underlying meso-vessels. Similarly, cortical thickness was evaluated to determine if greater tissue depth necessitates a higher density of vessel penetration to accommodate the expanded volume and spatial distribution of the branching hierarchy required to reach the capillary network. Finally, mean gray values were utilized as a control to identify potential algorithmic bias; a strong correlation here would indicate whether the detection sensitivity was confounded by local tissue intensity (e.g., preferential detection in darker, cell-dense regions) rather than true biological variation. See **Supplementary Figures 5-6** for two dimensional histograms showing the relation between meso-vessel density and other cortical metrics.

|  | Lobes | Nr. Vertices | Meso-Vessel Density vs Curvature |  | Meso-Vessel Density vs Thickness |  | Meso-Vessel Density vs Mean Gray Value (Merker) |  |
| --- | --- | --- | --- | --- | --- | --- | --- | --- |
| | | | Pearson ($r$) | Variance Explained ($r^2$) | Pearson ($r$) | Variance Explained ($r^2$) | Pearson ($r$) | Variance Explained ($r^2$) |
| **Left Hemisphere** | Frontal | 98,181 | -0.39 | 15.5% | -0.19 | 3.6% | -0.11 | 1.2% |
|  | Occipital | 51,112 | -0.33 | 11.1% | -0.21 | 4.3% | -0.18 | 3.1% |
|  | Temporal | 45,015 | -0.33 | 10.9% | -0.30 | 9.1% | 0.16 | 2.7% |
|  | Parietal | 44,351 | -0.43 | 18.9% | -0.23 | 5.2% | -0.12 | 1.5% |
|  | All Lobes | 238,659 | -0.37 | 14.0% | -0.26 | 6.5% | -0.08 | 0.6% |
| **Right Hemisphere** | Frontal | 87,288 | -0.40 | 16.1% | -0.24 | 5.7% | -0.11 | 0.4% |
|  | Occipital | 63,122 | -0.29 | 8.4% | -0.20 | 3.9% | -0.18 | 1.0% |
|  | Temporal | 40,492 | -0.41 | 16.8% | -0.35 | 12.2% | 0.16 | 3.8% |
|  | Parietal | 49,288 | -0.42 | 17.4% | -0.36 | 12.8% | -0.12 | 0.2% |
|  | All Lobes | 240,057 | -0.37 | 13.5% | -0.32 | 10.3% | -0.08 | 0.0% |

## 3 Discussion

Our maps reveal that intracortical vascular density is not uniform across the human brain at mesoscale. Instead, intracortical meso-vessel density varies substantially across the cortical landscape, following an organizational logic that frequently diverges from traditional neuronal boundaries. This suggests that the brain’s mesoscopic angioarchitecture possesses its own organizational logic, one likely driven by local metabolic budgets that are distinct from the structural properties of cytoarchitecture. While the functional implications remain to be explored, these maps provide a new anatomical measure that may help better understand the physiology of the active brain. In this context, the intracortical mesoscopic angioarchitecture might be viewed as an organizational feature, similar to the cytoarchitecture and myeloarchitecture features that have long guided our understanding of cortical structure.

### 3.1 Meso-vessel density: literature comparison

Our analysis identifies a mean meso-vessel density of approximately 0.7 vessels per mm^2^ (**Table 1**). This result is broadly consistent with established histological counts; for instance, Schmid et al. (2019) (synthesizing data from (Cassot et al., 2009; Lauwers et al., 2008)) reported 1.0 descending arteries and 0.5 ascending veins per mm^2^. While our estimate is lower than this combined total of 1.5 vessels per mm^2^, the values remain comparable given the methodological differences between study-specific protocols. A central finding of our work, however, is the angioarchitectural heterogeneity observed across the cortex at mesoscopic scale. This variability suggests that global averages, while useful as baselines, may not fully capture the distinct vascular topographies of different functional areas.

The variance in density estimates across the literature likely reflects differences in image resolution, partial volume effects, and the specific detection thresholds used to define “meso-vessels”. While the 100 micron isotropic resolution of the BigBrain dataset provides unprecedented coverage, it may naturally exclude the finest mesoscopic branches visible in more localized, higher-resolution preparations. For example, the landmark study by Duvernoy et al. (1981) reported a significantly higher density of 4.68 vessels per mm^2^ (see **Figure 6**). This higher figure likely reflects the unique advantages of Duvernoy’s specialized ink-injection preparations, which highlight the finest vascular ramifications, or may be influenced by localized sampling and histological shrinkage. Ultimately, these differences highlight the complementary nature of existing datasets and underscore the need for whole-brain mapping to establish the area-specific benchmarks required to understand regional variations in human cortical anatomy.

**Figure 6:**
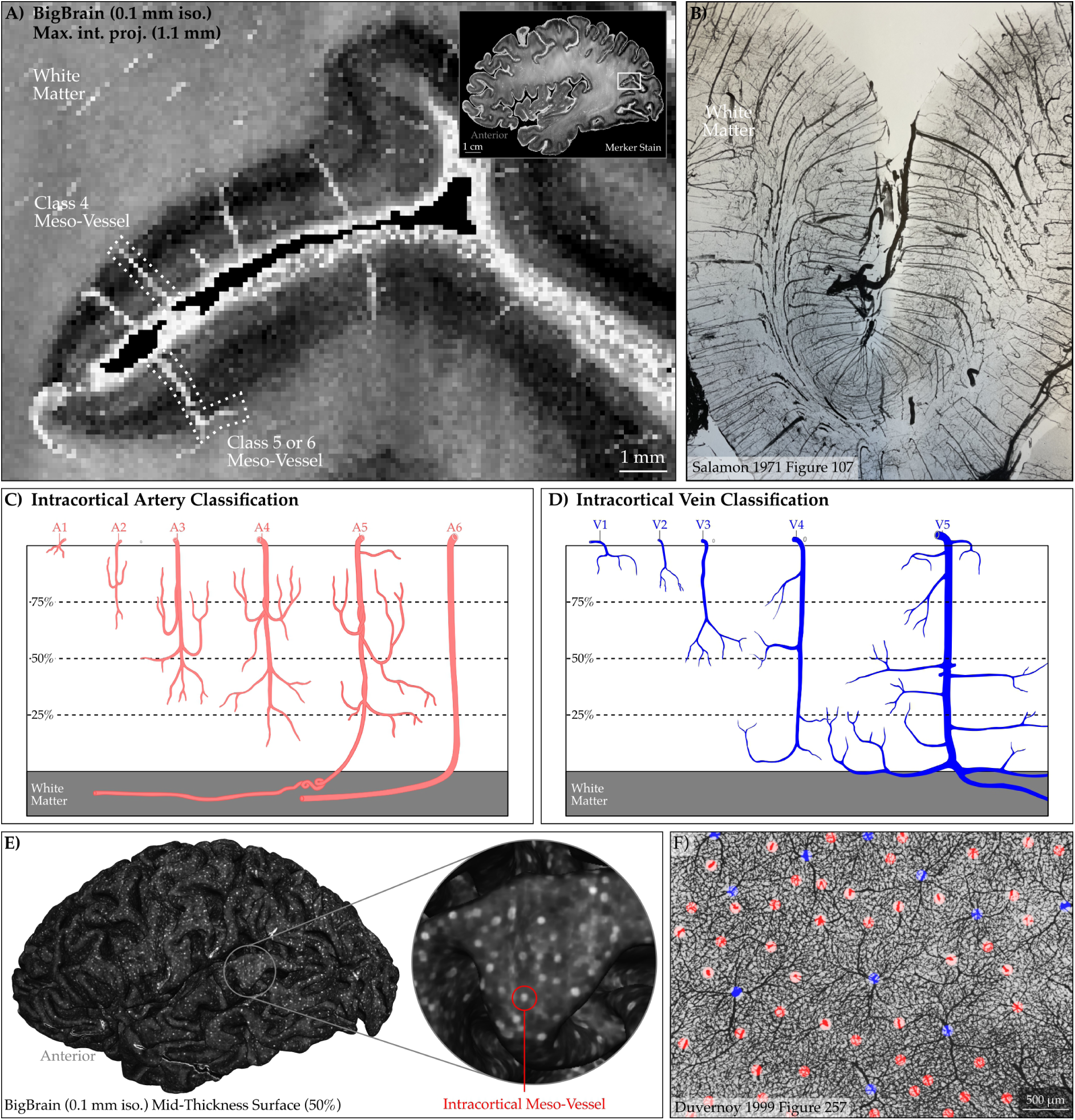
Validation of intracortical meso-vessels visible in BigBrain dataset. Here we show that the bright intracortical features visible within the cortex are intracortical meso-vessels. **Panel A** shows our meso-vessel projections in Merker-stained histology images. A sagittal 1.1 mm maximum intensity projection reveals the organized vertical lattice of the penetrating vascular network. **Panel B** shows an arterial microradiograms from Salamon (1971) (human brain). This figure does not have a scale bar; however the thickness of the slice is reported to be between 0.8 to 1.5 mm. Note that we are probably not seeing as many vessels in Panel A compared to Panel B because 100 micron isotropic resolution is not enough to capture the thinner vessels due to partial voluming. **Panels C-D** show intracortical artery and vein classifications, which are redrawn and rearranged based on Figure 248 of Duvernoy and Vannson (1999). Note that since we can not definitively know whether the vessels we see are arterial or venous vessels, we specifically use the term vessels. **Panel E** shows an entire hemisphere mid-thickness surface, highlighting meso-vessels as bright dots (see **Supplementary Figure 1** for visuals across cortical depths). **Panel F** shows a preparation by Duvernoy et al. (1981) which reveals the tangential view of the intracortical vessels following the physical removal (peeling) of superficial cortical layers. Not that we combined the Figure 63 and 64 from Duvernoy et al. (1981) which has the vessel segmentation together with Duvernoy and Vannson (1999) Figure 257 which has the scale bar.

### 3.2 Generalization of results beyond the BigBrain

While the BigBrain (Amunts et al., 2013) provides an unprecedented view of the mesoscopic network, it remains a map of a single human brain and hence cannot speak to diversity of vasculature characteristics across individuals. However, this dataset currently represents the highest-resolution record of an entire human brain in existence. While originally intended as a cytoarchitectonic reference (Paquola et al., 2021, 2025; Sitek et al., 2019; Wagstyl et al., 2018), we show that the BigBrain dataset contains a remarkably coherent, but so far neglected, vascular signal. It is the only whole-brain bridge between the macroscopic vessels found in medical textbooks and the microscopic capillaries studied in isolated slices.

While the BigBrain dataset provides a high-resolution anchor for human neuroanatomy, its hidden value lies in its power to calibrate the next generation of MRI imaging methods that can measure the vasculature in vivo. By quantifying the mesoscopic vascular architecture from 100 micron isotropic whole brain images, we provide the necessary blueprint to optimize and validate ultra-high-field MRI protocols, which excitingly can measure the intracortical mesoscopic vasculature in large numbers of humans with minimal effort (for macroscopic pial and leptomeningeal measurements see Bernier et al. (2018), Huck et al. (2019), and Mietzner et al. (2025)). Integrating our automated tracing image processing methods with advanced in vivo sequences, such as T_2_* weighted (Gulban et al., 2022, 2026; He et al., 2018) or time-of-flight (Bollmann et al., 2022; Haast et al., 2024; Xu et al., 2024) imaging, will allow the field to move beyond a single-subject atlas. We believe that this will transform mesoscopic vasculature from a histological curiosity to a critical measure of neurovascular health.

### 3.3 How intracortical meso-vessels might relate to other brain properties

Our maps reveal that the human brain’s mesoscopic vasculature follows a systematic logic governed by cortical geometry. The correlation between surface curvature and vessel density (r=-0.37 across both hemispheres, indicating sulci tend to have higher meso-vessel density) suggests that the physical folding of the cortical ribbon serves as a primary structural constraint, consistent with lower-resolution in vivo observations (Bernier et al., 2018). In this view, the mesoscopic vessels are not a stochastic distribution but a network that parallels the brain’s structural landmarks. In metabolically demanding areas, such as the primary somatosensory and visual cortices, intense energetic requirements likely exert a “biological pull” that overrides broad geometric trends (Keller et al., 2011; Weber et al., 2008).

On the other hand, the moderate strength of these correlations suggests that the vascular network is not merely a shadow of the brain’s physical volume. If the mesoscopic angioarchitecture was a simple function of space-filling, it would couple tightly with cortical thickness (r=-26 in left and r=-33 in right hemisphere); instead, its distribution might reflect a regional metabolic budget. In this view, the density of penetrating vessels represents a long-term structural investment in a region’s signaling capacity.

### 3.4 Technical limitations of the meso-vessel density map in this study

The meso-vessel density maps demonstrated in this study are subject to several technical constraints. While the Merker staining provides exceptional detail of the mesoscopic vascular trunks, it does not allow us to differentiate between penetrating arterioles and venules. In addition, there is no pial vasculature in this dataset; therefore we are strictly limited to the intracortical meso-vessels without their pial extensions that could have been informative for establishing their connections and branching.

Beyond staining constraints, the physical integrity of the BigBrain dataset introduces localized gaps in our vascular maps. In regions where the histological sections contain tears, folds, or tissue loss, meso-vessel density cannot be meaningfully sampled (see **Supplementary Figure 1**). While these artifacts are an inherent byproduct of large-scale histological processing, they underscore the necessity of expanding this framework to other imaging methods. Emerging mesoscopic in vivo human MRI acquisitions (Gulban et al., 2026; Serger et al., 2026; Stirnberg et al., 2024) that can cover the whole human brain within minutes at resolutions below 0.4 mm isotropic are going to be essential for overcoming such artifacts as well as allowing to scan more individuals more rapidly.

### 3.5 Future utility of our meso-vessel density maps

To facilitate adoption and cross-modal comparison, we have projected our whole-brain meso-vessel density maps onto the multi-modal cortical parcellation defined by Glasser et al. (2016). By anchoring these vascular metrics within this standard surface space, we provide a convenient reference for the neuroimaging community. Researchers can now directly correlate our vascular density values with existing datasets without the need for complex cross-modal registration. Our maps can be downloaded from https://doi.org/10.5281/zenodo.19954660.

## 4 Methods

### 4.1 Dataset

We utilized the BigBrain dataset (Amunts et al., 2012), a high-resolution (20 micron) histological reconstruction of a 65-year old post-mortem human brain. For our analyses, we utilized the “histological space, full volume, 100 micron with optical balancing” version under “3D Volume Data Release 2015” from https://bigbrain.loris.ca. We briefly refer to this dataset as “BigBrain 100 micron”. We choose to use BigBrain 100 micron to maintain a balance between mesoscopic detail and whole-brain computational tractability. **Figure 6** shows intracortical vessels directly visible in the BigBrain 100 micron dataset in comparison to other illustrations of the intracortical vessels captured in human brain slices.

### 4.2 Data Analysis

Our analyses are split into two main sections: meso vessel density mapping, and cortical surface visualizations.

### 4.3 Meso vessel density mapping

#### 4.3.1 Data size optimization

The BigBrain dataset at 100 micron isotropic resolution comprises nearly 2.6 billion voxels (2593392048, 1392 × 1541 × 1209) at 16-bit precision. To optimize computational performance and streamline downstream processing, we first constrained it to a field of view (FOV) encompassing only the cortex. This yielded approximately 1.8 billion voxels (1807800000, 1500 × 920 × 1310), nearly a 30 % reduction in data size (00 minc2nii bigbrain uint16.py)

#### 4.3.2 Cortical segmentation refinement

We first converted the existing cortical segmentations of the BigBrain dataset available as NIFTI images (Wagstyl et al., 2020). This segmentation indicates inner gray matter boundary, gray matter, and outer gray matter boundaries in three integers (see **Figure 2 Panel B**). To ensure histological-grade accuracy, we performed additional manual refinements of these segmentations using voxel-wise edits in ITK-SNAP v4.4.0 (Yushkevich et al., 2006). Specifically, we resolved “kissing gyri” instances where adjacent cortical banks are histologically distinct but voxel-wise contiguous to improve the precision of the pial and white-matter boundaries. Furthermore, we subdivided the hemispheres to allow our analyses to be run on each hemisphere separately.

#### 4.3.3 Whole brain voxel-wise computations

To establish a biologically grounded coordinate system for cortical depth dimension, we utilized the LN2 LAYERS program from the LayNii (v2.9.0) (Huber et al., 2021). This allowed for the voxel-wise computation of equi-volume cortical depth, curvature, and thickness across the entire hemispheric manifold (01 run LN2 LAYERS.py).

To manage the high computational demands of the subsequent vessel-tracing algorithms, we parcellated each hemisphere into 250 cortical cells. These parcels cells generated using a two-step process on the whole brain images:

1. Sampling: We identified a set of seed points across the hemispheric surface using an iterative farthest points algorithm (LN2 IFPOINTS program), ensuring approximately equally distant coverage (02 run LN2 IFPOINTS.py).
2. Parcellation: These 250 seeds served as the centroids for a Voronoi diagramming algorithm (`LN2 VORONOI` program), which subdivided the cortical geometry into computationally manageable, mutually exclusive and collectively exhaustive cells, which can be seen in **Figure 2 Panel D** (03 run LN2 VORONOI.py).

#### 4.3.4 Chunk-by-chunk voxel-wise computations

1. **Chunking for computational efficiency:** After parcellating the brain, we divided the 100 m isotropic images into manageable chunks. We centered each chunk on a specific cell centroid within a 384 × 384 × 384 voxel window, yielding 250 chunks per hemisphere (see **Figure 2 Panel G**). This modularity allowed us to process high-resolution data within standard computing memory limits in the next steps. We applied this chunking to the Merker stain values, cortical segmentations and other metrics computed in the whole brain steps (01 chunk cells.py, 02 chunk anat.py, 03 chunk rim.py, 04 chunk midgm.py, 05 chunk points.py, 06 chunk metric equivol.py).
2. **Enhance vessel contrast:** To bring out the subtle traces of intracortical vessels in the Nissl data, we first applied one step of grayscale dilation (replacing the central voxel with the maximum value encountered within the 3 × 3 × 3 kernel). This expanded the bright, thin vessel traces, making them more prominent against the gray matter. We then calculated the gradient magnitude (square root of the sum of the squares of each spatial gradient, (Gulban et al., 2018)) to sharpen the vessel borders. Finally, a grayscale closing operation (one-step dilation followed by one-step erosion) filled in the gaps within the intracortical vessel trunks (07 enhance vessels.py).
3. **UV Parametrization and 3D Flattening:** To map the cortex, we established an orthogonal coordinate system on the mid-thickness voxels using the LN2 MULTILATERATE program from LayNii (as established in (Gulban et al., 2022)). For each cell centroid, we computed UV coordinates indicating directions tangential to the cortical surface and a D coordinate indicating equivolume cortical depth. Each resulting UV geodesic disk had a 17 mm radius. It is important to note that we computed the intrinsic coordinate system for each cell separately. This localized approach minimizes the geometric distortion that would occur if we attempted to flatten the entire cortex at once, a strategy analogous to the cuts used in UV mapping methods to prevent mesh stretching (see Botsch et al. (2010) for further details). We then used the UVD coordinates to flatten each cortical chunk, transforming the data from the cardinal XYZ axes (anterior-posterior, inferior-superior, left-right) into an intrinsic, cortex-referenced UVD axes (Gulban & Huber, 2024). To ensure the fine intracortical details remained intact during this projection, each flat patch was sampled at a resolution of 1000 × 1000 voxels (U and V) by 150 voxels (D). This high-density grid prevents the smoothing or mixing of features that lower resolution grids might cause, preparing the data for automated vessel detection (08 make control points.py, 09 run LN2 MULTILATERATE on chunks.py, 10 run LN2 PATCH FLATTEN on chunks.py).
4. **Detecting vessels:** We identified vascular candidates by applying a 2D local maximum operator to each slice of the flattened 3D stacks. To bridge small gaps in the traces caused by histological artifacts, we applied a 2D morphological dilation to the detected points. We then filtered out transient noise by measuring cross-laminar persistence. This metric (similar to the “columnarity index” used in Pizzuti et al. (2023)) tracks how consistently a vessel point appears through the depth-wise layers of the stack; we retained only those vessels that persisted through more than 40 % of the layers (11 detect vessels.py). The result of this processing was a 2D binary image of vessels. We then assigned a unique integer to each distinct vessel using a connected cluster labeling algorithm (LN2 CONNECTED CLUSTERS)
5. **Compute meso-vessel density:** To quantify the vessel distribution, we calculated the meso-vessel density across each patch. We slid a circular kernel (90-pixel radius, 3 mm in diameter as measured from on the mid-thickness voxels geodesically) across the labeled image to count the number of unique vessels within the window (12 compress to 2D.py, 13 connected clusters.py, 14 count vessels.py). Then we divided each measurement with the counting kernel area (7.07 mm^2^) multipled by the areal shrinkage correction factor (1.55 as taken from Wagstyl et al. (2020)) to transform the counts into “meso-vessel den-sity” measured per unit mm^2^. Critically, because this density estimation is performed in absolute units of distance within a volumetric patch, rather than being calculated per vertex on a surface mesh, our results are not subject to the curvature biases associated with varying vertex density as shown in Kay et al. (2019) Figure 2 Panel C. An example map of a single patch can be seen in our **Figure 2 Panel H**.
6. **Back projection:** After calculating the meso-vessel densities, we returned the data to its original anatomical context. We unflattened the patches by back-projecting the voxels from the UVD system into the convoluted brain (XYZ) using the LN2 PATCH UNFLATTEN program (15 decompress to 3D.py, 16 run LN2 PATCH UNFLATTEN on chunks.py). Finally, we stitched these individual chunks back into the whole-brain reference frame (17 unchunk.py). Where patches overlapped, we used a voxel-wise maximum operator to merge the values. We preferred the maximum operator to eliminate low density edge artifacts that occur at the flat patch edges. We ensured substantial overlap between the chunks to prevent edge artifacts from distorting the density measurements. This step results in a whole brain image that has a meso-vessel density value for each cortical gray matter voxel.

### 4.4 Cortical surface visualizations

#### 4.4.1 Reconstruction of laminar meshes

We visualized intracortical vessels through a multi-step triangular mesh reconstruction designed to preserve fine-scale vascular structures across the cortex. To accurately represent the laminar architecture, we first generated four equidistant geometric layers using LN2 LAYERS program in LayNii. From these, three triangular mesh surfaces were reconstructed at 25 %, 50 %, and 75 % cortical depth using the marching cubes algorithm in BrainVoyager.

Unlike conventional fixed-vertex methods we utilized varying vertex density meshes. This is done to prevent insufficient sampling of superficial layers where a single mesh is projected outward. Our varying vertex density meshes ensure that the increasing surface area from deep to superficial layers is fully represented. For instance, in the right hemisphere, vertex counts increased from the deep layer with 12499138 vertices, to the middle layer with 12923304 vertices, and to the superficial layer with 13365118 vertices. Note that our varying vertex density approach to triangular mesh sampling is critical for conveniently visualizing intracortical meso-vessels without using more resource intensive sampling methods such as upsampled fixed-vertex methods (Polimeni et al., 2018), or texture wrapping (Botsch et al., 2010; Gao et al., 2015).

Following reconstruction, meshes were refined using advanced mesh smoothing (150 iterations, 0.07 force) to eliminate tessellation artifacts while preventing shrinkage. Voxel values were then sampled exactly at mesh vertices via BrainVoyager v24.2.5 (Goebel, 2012) “Create SMP (Surface Map)” function, a strategy chosen specifically to avoid spatial blurring and preserve the high-resolution details of the intracortical veins. Finally, to ensure visibility within sulcal regions, surfaces were inflated (32000 steps) and rendered with a low shininess parameter (0) to provide a subtle mesh lighting for clear visual assessment.

#### 4.4.2 Mid-thickness mesh processing

We projected the meso-vessel density maps onto reconstructed cortical mid-thcikness surfaces to compare their distribution with other structural metrics, such as curvature, thickness, and Merker-derived gray values. To make the surfaces easier to load and navigate, we simplified the mesh geometry using advanced mesh smoothing (with vertex displacement contrast at 150 iterations) and vertex decimation (250000 vertices). This smoothing and downsampling did not detract the quality of our visualizations as the relevant maps were already smooth quantities that were naturally computed beyond single voxel resolution. The resulting triangular mesh surface can be seen in **Figure 2 Panel C**.

We then sampled the voxel data onto the resulting surface vertices using “sample volume data exactly at mesh vertices”, which can be seen in **Figure 2 Panel F**. To ensure a clean visualization, we manually masked the corpus callosum, (which is severed during hemispheric separation) to exclude non-cortical values from the final maps. While these surface-based steps allow for global comparison and visualization, the underlying data remains anchored in the original voxel space.

#### 4.4.3 Registration to HCP MMP1.0 meshes

We aligned the BigBrain mid-thickness surface to the (Glasser et al., 2016) surface atlas using Cortex-Based Alignment (CBA) (Frost & Goebel, 2012, 2013; Gulban et al., 2020) as implemented in BrainVoyager. To ensure a robust alignment, the BigBrain mid-thickness surface was mildly inflated prior to computing cortical curvature. This step was necessary to harmonize the high-frequency topological details of an individual histological reconstruction with the smoother curvature profile characteristic of the HCP MMP1.0 mesh. By calculating curvature on a mildly inflated mesh, we generated a feature map with a spatial frequency comparable to the atlas, facilitating more accurate vertex-to-vertex matching.

Following this preprocessing, we employed the standard CBA “align pair” protocol to establish a discrete surface-to-surface correspondence between the HCP MMP1.0 and the Big-Brain mid-thickness mesh. This mapping allowed us to project our mesoscopic vessel density measurements directly onto the Glasser atlas for regional analysis (as visualized in **Figure 4** and **Supplementary Figures 2-4**). To facilitate broad utility, our mesoscopic vessel density maps are provided for both hemispheres of the Glasser atlas across a multi-resolution framework, ranging from 32K to 128K vertices per mesh.

#### 4.4.4 Correlation analysis across mesh vertices

To assess the consistency of vascular-structural relationships across the cortex, we partitioned the hemispheric triangular mesh surfaces into its four principal lobes: frontal, parietal, temporal, and occipital. In addition, note that the corpus callosum region was excluded. The boundaries were manually defined by an expert directly on the triangular mesh surfaces according to gross anatomical landmarks (O.F.G.) using surface drawing tools as implemented in BrainVoyager mesh viewing window. This coarse parcellation allows us to assess whether the observed correlation patterns remain stable across the broader cortical landscape.

For the vertices within each lobe, we calculated Pearson correlation coefficients (r). These comparisons, detailed in **Table 2**, allow us to determine the degree to which the density of the mesoscopic vascular network aligns with the geometric and cytoarchitectonic features of the brain.

## Acknowledgements

O.F.G. and R.G. have financial interest tied to Brain innovation. The authors thank Robert Turner for his valuable comments and insights on an early version of the preprint.

## Supplementary Material

**Supplementary Figure 1:**
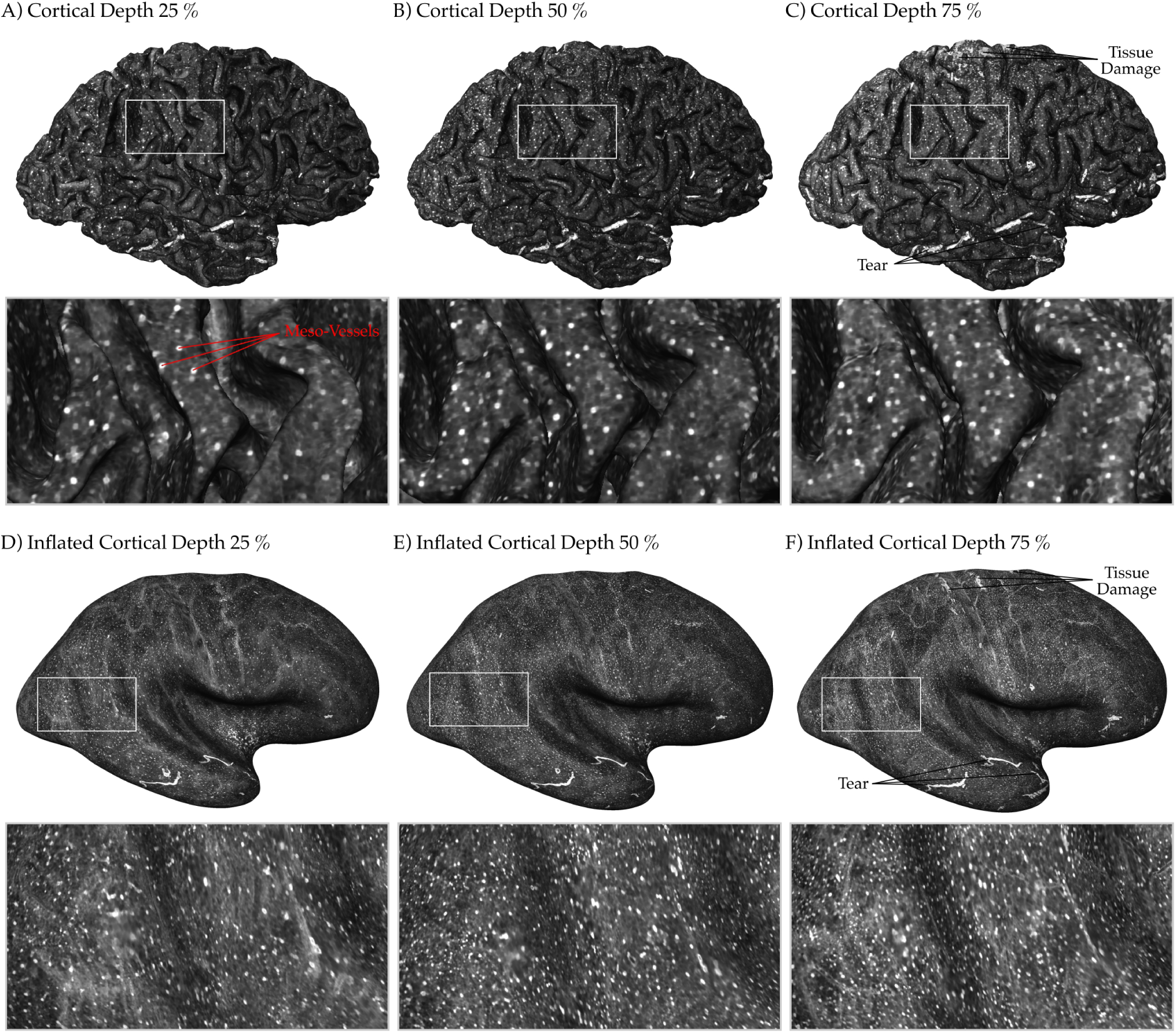
Intracortical meso-vessels are visible across cortical depths of the BigBrain dataset. **Panels A-C1** show triangular mesh reconstructions at three thickness-normalized (equidistant) geometric layers. The intracortical meso-vessels are visible as distinct, high-intensity circular dots distributed across the cortical landscape in the vessel enhanced contrast. **Panels D-F** show corresponding inflated surfaces, which reveal meso-vascular structures previously hidden within the sulci. Collectively, these visualizations demonstrate an increase in the density of visible meso-vessels from deep toward superficial cortical layers (more bright dots in superficial layer than the deep layer). This depth-dependent distribution aligns with the classical anatomical observations of Duvernoy et al. (1981) where there are more numerous but smaller radial vessels in the superficial layers of the cortex. Localized tissue damage resulting from histological processing appears as larger high-intensity clusters.

**Supplementary Figure 2:**
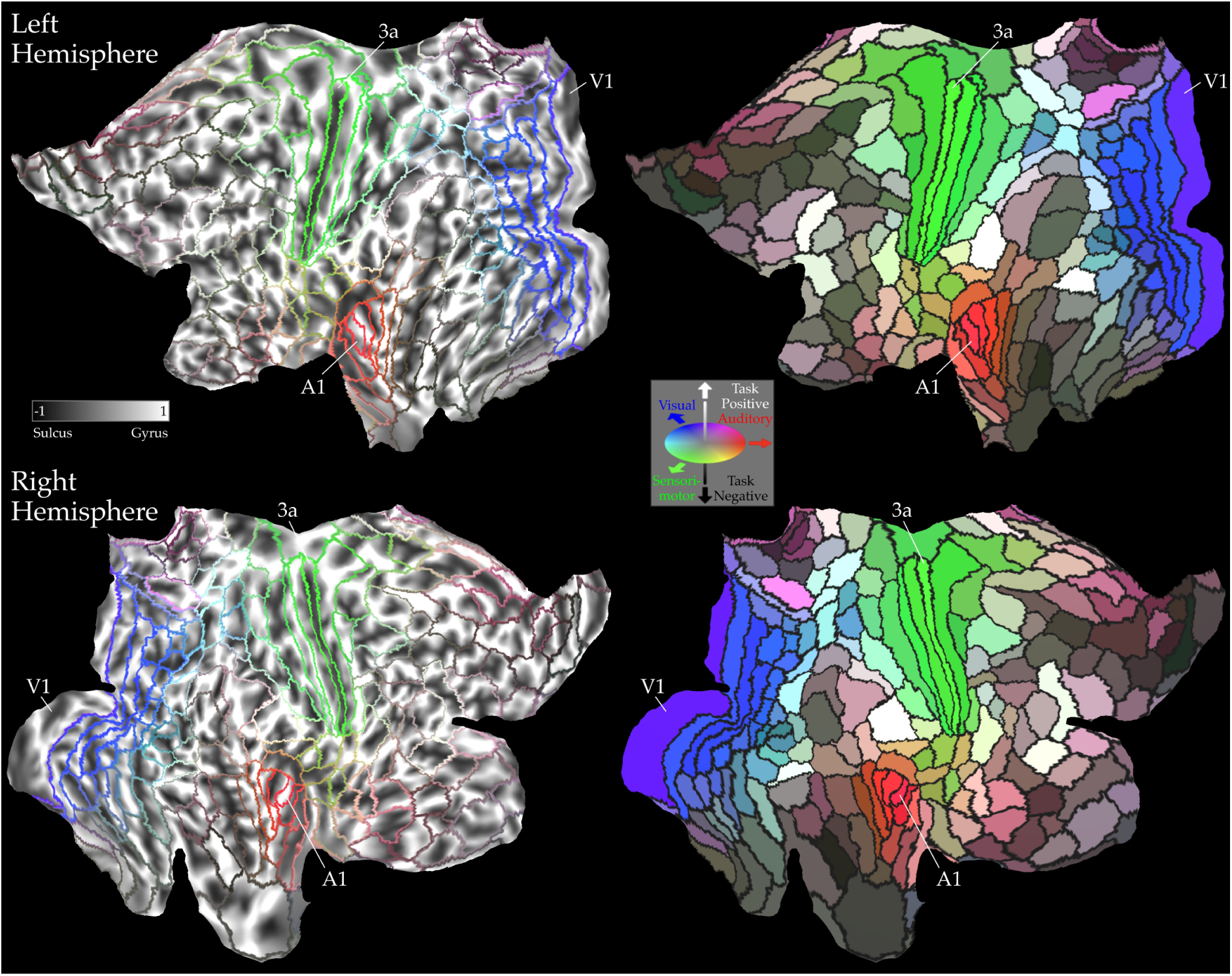
Cortical curvature maps projected onto flattened cortical surfaces of the HCP’s multimodal cortical parcellation (HCP MMP1.0) from Glasser et al. (2016).

**Supplementary Figure 3:**
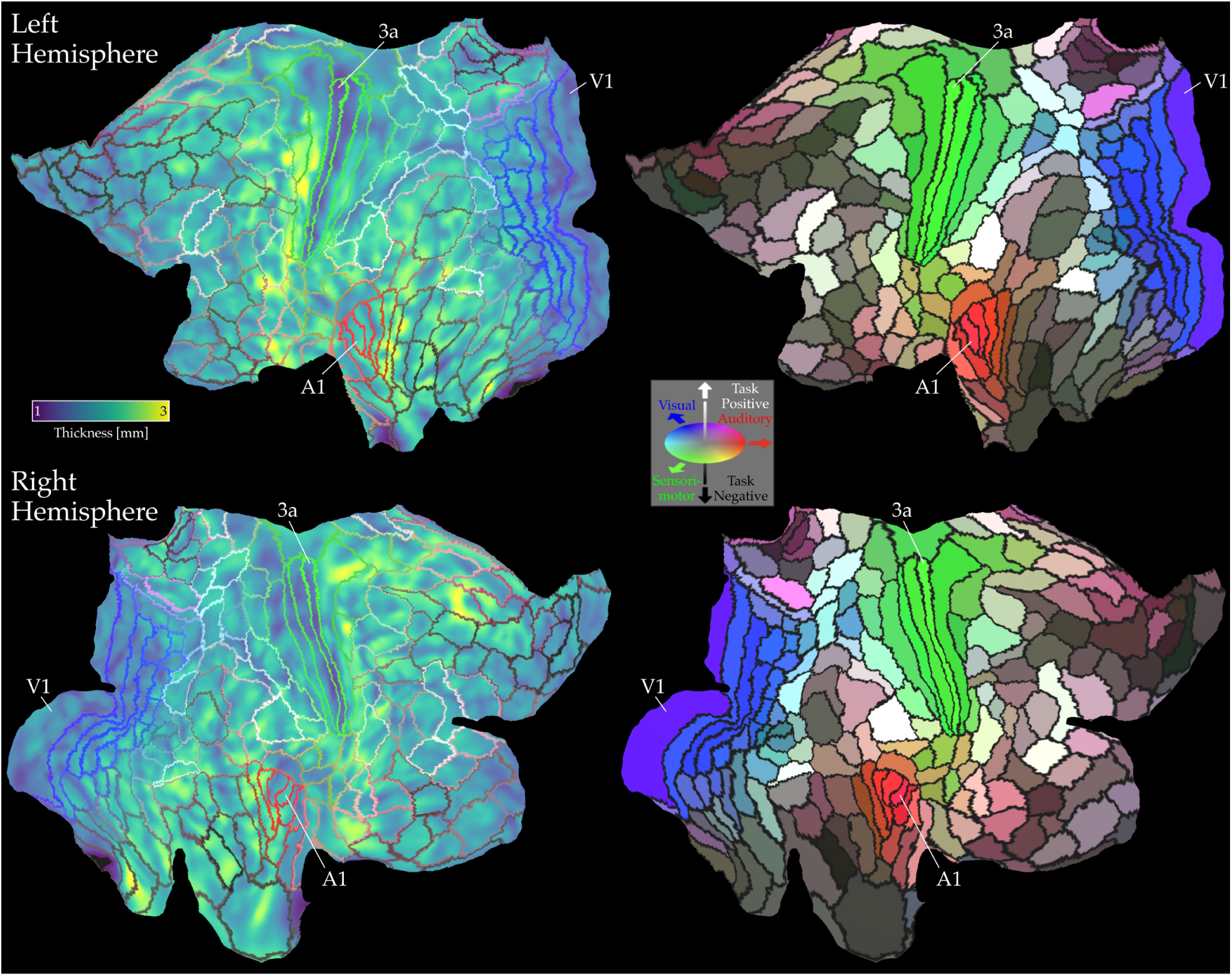
Cortical thickness maps projected onto flattened cortical surfaces of the HCP’s multimodal cortical parcellation (HCP MMP1.0) from Glasser et al. (2016).

**Supplementary Figure 4:**
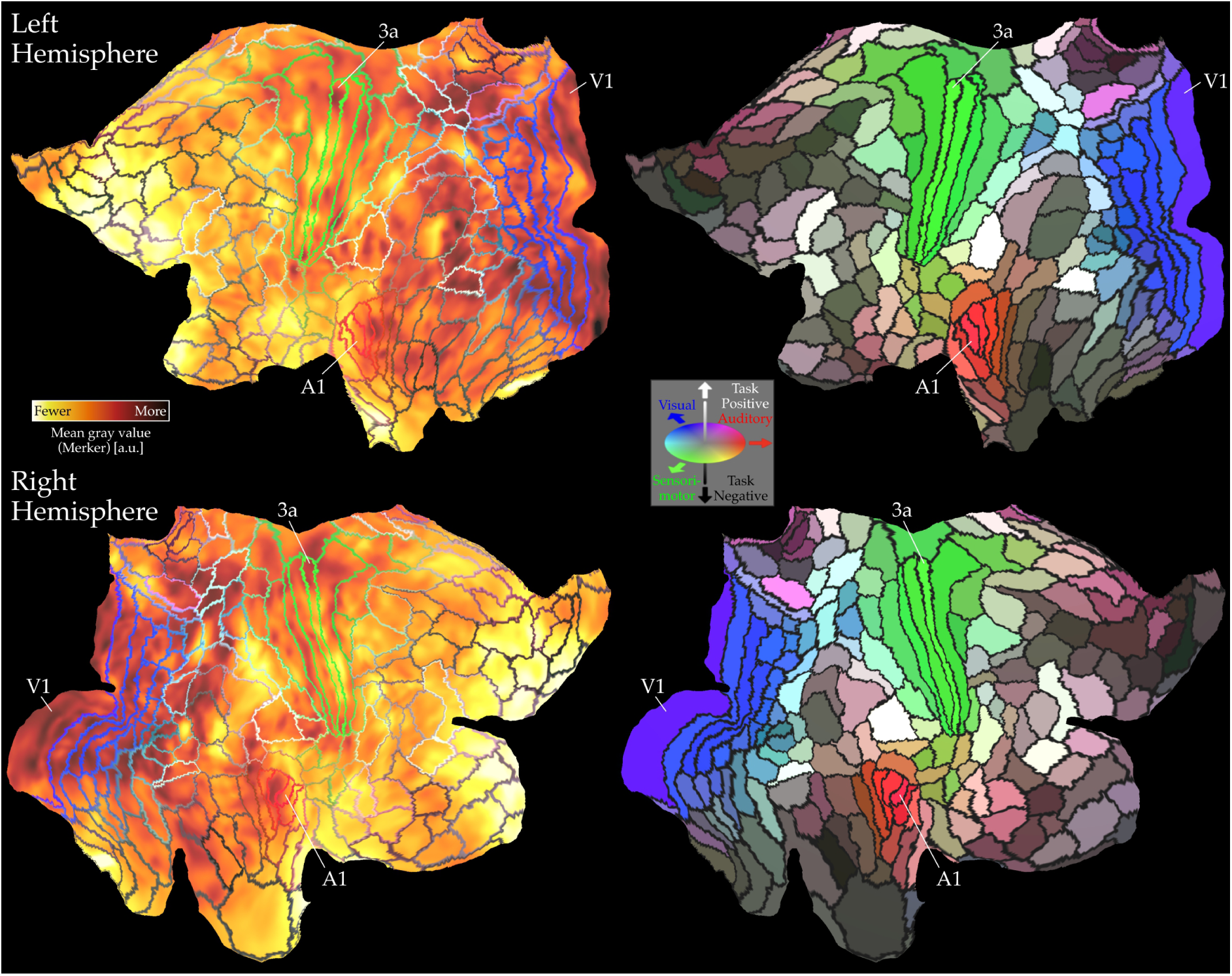
Mean gray value (Merker) maps projected onto flattened cortical surfaces of the HCP’s multimodal cortical parcellation (HCP MMP1.0) from Glasser et al. (2016)

**Supplementary Figure 5:**
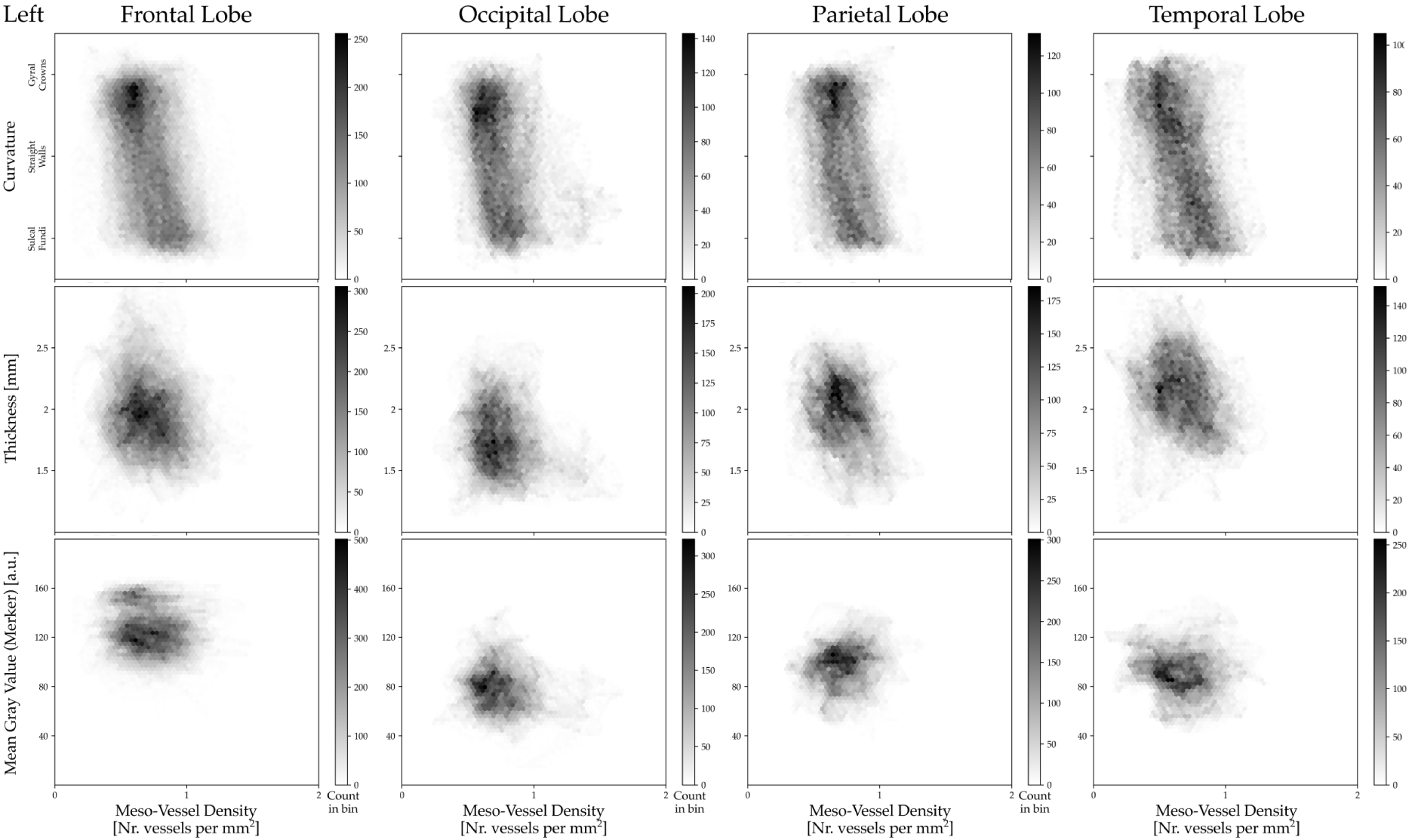
Two dimensional histograms showing the relation between meso-vessel density and other cortical metrics for the left hemisphere across the vertices of the triangular surface mesh. Qualitatively, it seems that, between hemispheres, curvature correlation is higher in frontal and parietal, lower in occipital, and less consistent in temporal. The thickness correlation is highest in temporal.

**Supplementary Figure 6:**
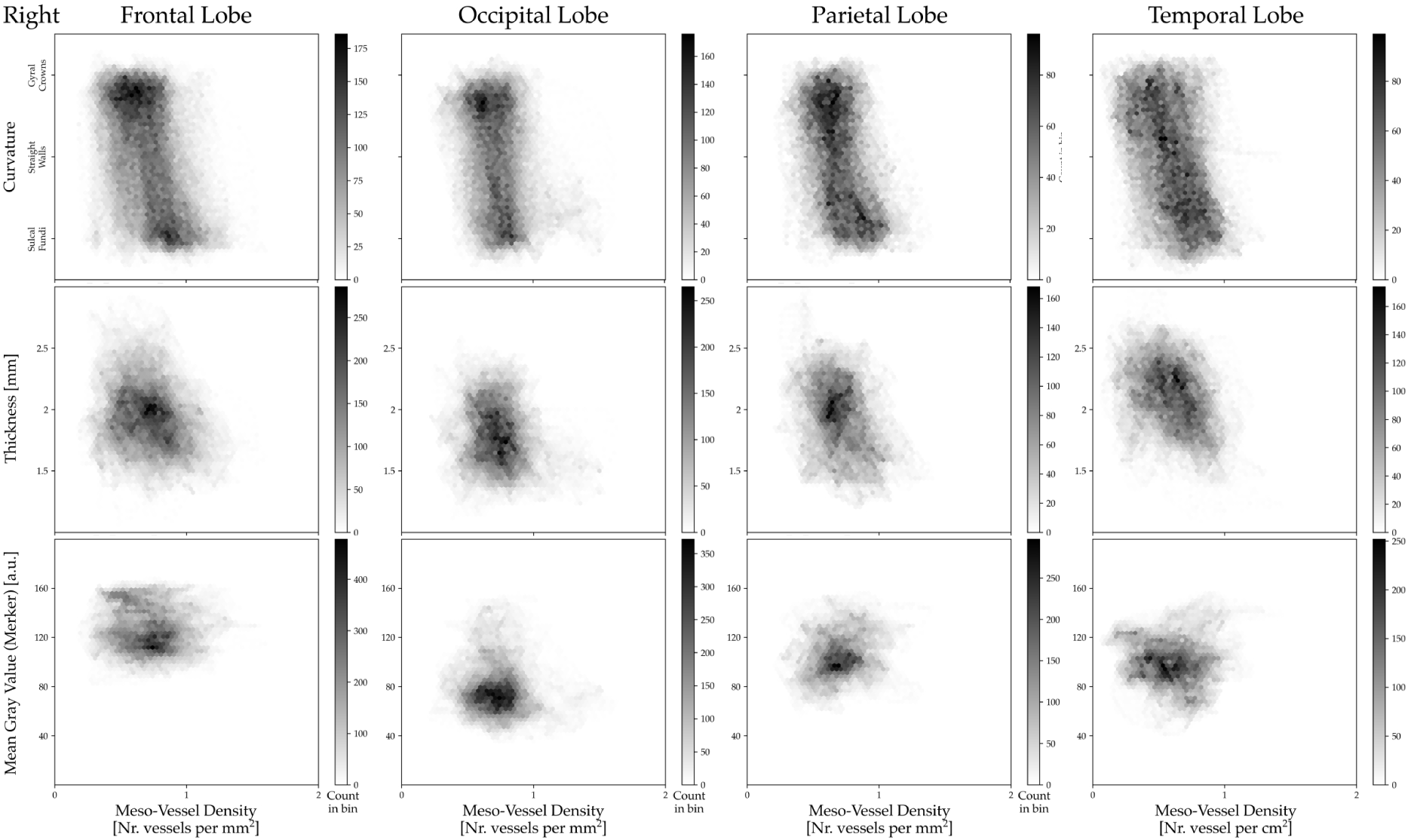
Same as Supplementary Figure 4 but plots show the data from the right hemisphere.

## Notes

https://doi.org/10.5281/zenodo.19954660

### Competing Interest Statement

The authors have declared no competing interest.

### Summary of Updates

Minor fixes grammar and typos. And a few new citations are added.

